# Host brain environmental influences on transplanted medial ganglionic eminence progenitors

**DOI:** 10.1101/2024.01.15.575686

**Authors:** Rosalia Paterno, Thy Vu, Caroline Hsieh, Scott C. Baraban

## Abstract

Interneuron progenitor transplantation can ameliorate disease symptoms in a variety of neurological disorders. This strategy is based on transplantation of embryonic medial ganglionic eminence (MGE) progenitors. Elucidating host brain environment influences on interneuron progenitors as they integrate is critical to optimizing this strategy across different disease states. Here, we systematically evaluated age and brain region influences on survival, migration and differentiation of transplant-derived cells. We find that early postnatal MGE transplantation yields superior survival and more extensive migratory capabilities compared to juvenile or adult. MGE progenitors migrate more widely in cortex compared to hippocampus. Maturation to interneuron subtypes is regulated by age and brain region. MGE progenitors transplanted into dentate gyrus sub-region of early postnatal hippocampus can differentiate into astrocytes. Our results suggest that host brain environment critically regulates survival, spatial distribution and maturation of MGE-derived interneurons following transplantation. These findings inform and enable optimal conditions for interneuron transplant therapies.

## Introduction

Cell transplantation is a promising therapy for neurological disease as it provides an opportunity to modify neuronal networks. Debilitating conditions, such as epilepsy, neuropsychiatric disorders, Alzheimer’s disease or schizophrenia are characterized by an inhibitory synaptic transmission alteration linked to loss of sub-populations of GABAergic interneurons ^1–5^. These pathologies, collectively referred to as “inter-neuropathies” ^6^, profoundly alter neuronal networks contributing to generation of abnormal brain activity. Neurostimulation approaches, such as deep brain stimulation (DBS) or responsive neurostimulation (RMS), provide a degree of control by directly stimulating inhibitory pathways within dysfunctional brain networks ^7,8^. These approaches require intracranial placement of electrodes in targeted brain regions, but do not provide a selective activation of inhibition as local excitatory neurons are also stimulated. A cell transplantation strategy capable of introducing subsets of inhibitory interneurons to selectively modify GABAergic circuits, in specific brain regions implicated in these diseases, offers a more targeted network-based option. However, to be effective, a cell transplantation therapy should satisfy some critical conditions: (i) adequate cell survival in host brain following transplantation, (ii) effective migration in targeted host brain regions, (iii) differentiation into specified interneuron sub-types and (iv) functional integration into existing neuronal networks. Embryonic medial ganglionic eminence (MGE) progenitor cells appear to satisfy these criteria ^9^. Indeed, embryonic MGE cell transplantation has shown promising results in preclinical models representing several of these neurological diseases^10–13^.

The overall premise for using embryonic MGE progenitors for cell transplantation was initially established by Wichterle and colleagues ^14^. Using MGE progenitors from mouse embryos they demonstrated that these cells give rise to GABAergic interneurons following transplantation into cortex of host pups at postnatal day 2 (P2). MGE is a transient embryonic structure defined anatomically and by its unique transcription factor profile ^15,16^. Progenitor cells from this sub-pallial proliferative zone migrate tangentially in streams to populate neocortex, hippocampus and striatum. Upon reaching these brain regions, MGE progenitors differentiate into two major GABAergic interneuron sub-classes: (1) parvalbumin-expressing (PV) fast-spiking interneurons, including basket and chandelier cells and (2) somatostatin-expressing (SST) interneurons mostly represented by Martinotti cells ^17^. Similar to normal development, transplanted MGE progenitors exhibit robust migratory and differentiation capabilities ^18^. Moreover, from a network perspective, MGE-derived interneurons functionally integrate in host brain where they receive synaptic input from endogenous neurons, make appropriate layer-specific synaptic connections onto endogenous excitatory neurons ^19,20^ and enhance GABA-mediated inhibition ^21^ in a sub-type specific manner i.e., PV-positive MGE-derived neurons innervate cell somas and SST-positive MGE-derived interneurons innervate dendritic regions.

Despite the success of embryonic MGE progenitor transplantation to ameliorate disease symptoms in animal models of Alzheimer’s disease ^11,13^, Parkinson’s disease ^22^, epilepsy ^10,23^, neuropathic pain ^24^, traumatic brain injury (TBI) ^12^, schizophrenia ^25^, and autism ^26^, optimization of this approach for specific brain regions and developmental stages has not been addressed. Although not directly explored in prior studies, as yet unknown aspects of the host brain environment are likely to be critical to effective transplantation and integration of MGE-derived interneurons for such a wide variety of neurological disorders. One notable issue among prior adaptations of MGE transplantation is the reported variability of MGE-derived cell survival which ranges from 1% to 20% ^18,27,28^. A second issue is the number of injections necessary to obtain adequate migration of cells across the targeted brain region with some publications reporting cell concentrations as high as 200,000 cells per injection and others only 15,000 per site ^21,29^. A third issue is developmental stage of the recipient animal as limited migration and parvalbumin-positive cell differentiation were observed in some adult transplants. Taken together, these issues suggest that host brain environment is a critical determinant in the survival, migration and differentiation of MGE-derived interneurons following transplantation.

Here we address these issues to better understand environmental factors and conditions influencing effective integration of embryonic MGE progenitors following transplantation. Embryonic MGE progenitors were transplanted during different developmental stages (pup, juvenile, and adult) and into different brain regions critical to Alzheimer’s, epilepsy, TBI, schizophrenia and autism (cortex and hippocampus). Sacrificing recipient animals at 30 days after transplantation (DAT), we carefully assessed survival, migration, and maturation profiles. Micro-targeting of MGE cell transplantation to specific hippocampal sub-regions was also explored in an effort to potentially tailor this strategy for more focal manipulation of neuronal circuitry. Early postnatal transplantation yielded better MGE-derived cell survival rates and more extensive migratory capabilities compared to juvenile and adult brain transplantation. Host brain region also influenced these factors with cortex showing a greater capacity to receive MGE-derived interneurons than hippocampus. Within hippocampus, transplanted MGE progenitors showed unique migration and differentiation fates depending upon hippocampal sub-region where MGE progenitor cells were initially deposited. Altogether, this information highlights age- and region-specific environmental factors that influence successful implementation of embryonic MGE progenitor cell transplantation as a potential therapy for neurological diseases.

## Material and Methods

### Animals

All experiments were performed on mice maintained on a 12-hour light/12-hour dark cycle with no food or water restrictions. All procedures involving animals followed guidelines of the National Institutes of Health and were approved by Institutional Animal Care and Use Committee at University of California, San Francisco (#AN181254-02B). Embryonic donor tissue was obtained by crossing wild-type CD1 mice (Charles River Laboratories, Cat Num #022) to homozygous beta-actin EGFP mice on a CD1 background. We used male and female CD1 and male C57BL/6J (Jackson laboratory; 000664) mice as recipients, as specified in each experiment. The study was conducted in accordance with the ARRIVE (Animal Research: Reporting of in Vivo Experiments) guidelines.

### Tissue dissection and transplantation

MGE progenitor cells were harvested from E13.5 GFP+ transgenic embryos as previously described ^10,13^. In brief, embryonic day 0.5 was defined when the vaginal plug was detected. To dissect MGE, embryonic brains were first removed, telencephalon isolated and the two hemispheres separated by a sagittal cut. The ventral telencephalon was exposed, and medial ganglionic eminence was isolated using the sulcus that clearly divide the medial from lateral ganglionic eminence. The dorsal MGE was collected after removing the preoptic area and mantle zone. MGE progenitor cells were collected in Leibovitz L-15 media, mechanically dissociated in a single cell suspension in media containing DNase I (Roche, 100 ug/ml) using repeated pipetting and concentrated by centrifugation (3 min at 800 X g). Cells were kept at 4° C until transplantation.

Concentrated GFP+ MGE cells (∼600 cells per nl) were injected using 30° beveled glass micropipettes (80 μm diameter tip, Witerol 5 μl, Drummond Scientific) prefilled with mineral oil into cortex or hippocampus of recipient pup (P2-4), juvenile (P30-40) and adult (P90-150) animals. Pipettes were attached to a microinjector (Narishige) mounted to a stereotaxic machine (Kopf). We quantified cell viability of MGE cells using Triptan Blue (Sigma) and a viability more than 80% was considered as cutoff to proceed with transplantation. Each cortical recipient animal received a single injection containing 5 x 10^4^ cells. Hippocampal recipient animals received a single or a double injection per hemisphere containing 3 x 10^4^ cells due to increased risk of clumping when used higher volumes.

Pups were anesthetized by exposure to ice until pedal reflexes were absent. Around 50-100 nl of highly concentrated cells were injected at the following coordinates from bregma to target cortex (1.2 mm anterior, 1.2 mm lateral and 0.6 mm dorsal) or hippocampus (1.2 mm anterior, 1.2 mm lateral and 1.4 mm dorsal). After injection pups were returned to their mother. Juvenile and adult animals were anesthetized by a mixture of ketamine/xylazine. Cell injections were made into cortex at the following coordinates form bregma: anterior-posterior (AP) – 1.75 mm, medial lateral (ML) 1.75 mm, dorsal ventral (DV) 0.6. For hippocampus to target radiatum layer of area CA3 or stratum oriens of area CA1 were made at the following coordinates: AP – 2 mm, ML 2.5, DV 1.8 and AP – 2 mm, ML 1.6, DV 1.15 respectively.

### Immunohistochemistry

Mice were deeply anesthetized with a mixture of Ketamine/Xylazine and perfused transcardially with saline followed by 4% paraformaldehyde (PFA; Electron Microscopy Science). Brains were post-fixed with PFA 4% overnight and 50 µm thick coronal sections were cut with a VT 1000S vibratome (Leica Microsystems Inc., Buffalo Grove, IL). Free-floating vibratome sections were processed as follows. Sections were washed in PBS and blocked for 1 h in PBS containing 10% goat serum at room temperature. Sections were then incubated overnight in primary antibodies at 4° C and then washed in PBS and incubated with secondary antibody for 2 h at room temperature in the dark. Primary antibody used: anti-NeuN (Millipore; MAB377; 1:500), anti-GFP (AvesLabs; GFP-1020; 1:1000), anti-parvalbumin (Sigma-Aldrich; P3088; 1:1000), anti-somatostatin (Santa Cruz Biotech, SC55565, 1:200), anti-GABA (Sigma, A2052, 1:500), anti-GFAP (Millipore, MAB3402, 1:500), anti-Oligo2 (Millipore, AB9610, 1:300), anti-CC1 (Sigma-Aldrich; OP80; 1:500). Secondary antibody used: Alexa 488 (Invitrogen; A11039; 1:1000), Alexa 594 (Invitrogen; A11005; 1:1000) and Alexa 647 (Invitrogen; A21244; 1:1000).

### Cell counts and quantification

Images were acquired at 1024 pixels resolution using a Nikon confocal microscope. Quantification analysis was performed using NIS-Elements (Nikon software) on fluorescent labeled sections (50 µm) imaged with 4X, 10X and 40X objectives. All transplanted cells that expressed GFP were counted in every sixth coronal section (300 µm apart) in all layers of cortex or hippocampus, as previously described ^21^. To estimate cell survival in each hemisphere, we counted the number of GFP+ cells in every 6-coronal section (300 µm apart) through the rostro-caudal axis and multiply this number by 6, we then calculated then number of (total survived cells/numbers of transplanted cells) *100. To assess distribution of GFP+ cell across cortical layers, GFP+ cells were allocated to one of 10 bins superimposed on the area between pial surface (bin 1) and white matter (bin 10). To assess the percentage of grafted GFP+ cells co-expressing NeuN, SST, PV, GFAP, Oligo or CC1 after transplantation (n = 3 mice per marker), we quantified the number of GFP+ cells co-expressing these antibody markers in every sixth coronal section (300 µm apart).

### Statistical analysis

All analyses were performed using PRISM (GraphPad). To compare different groups, we used two-tailed t-tests, unpaired t-tests and one-way ANOVA’s as specified in each analysis. All plots with error bars are reported as mean ± SEM. No data were excluded from analysis.

### Data availability

The original data of this study are available from R.P. upon reasonable request.

## Results

Murine embryonic MGE progenitor cells migrate widely following early postnatal transplantation into recipient mouse brain ^18^. Within host brain, transplanted MGE-derived interneurons efficiently distribute across cortical layers ^30–32^. However, residual clustering and limited migration away from the injection site have also been reported ^29,33^. Since factors transiently expressed in developing brain e.g., clustered gamma-protocadherins ^34,35^ or CCCTC-binding factor ^36^ can influence migration and integration of MGE progenitor cells, we here investigated integration following transplantation into recipient mice at different developmental ages. Because factors expressed in local microcircuits e.g., vesicular GABA transporters ^32^ or MTG8 ^37^ can also influence integration of these progenitors, we compared transplantations into cortex and hippocampus (Fig. 1A).

**Figure 1:**
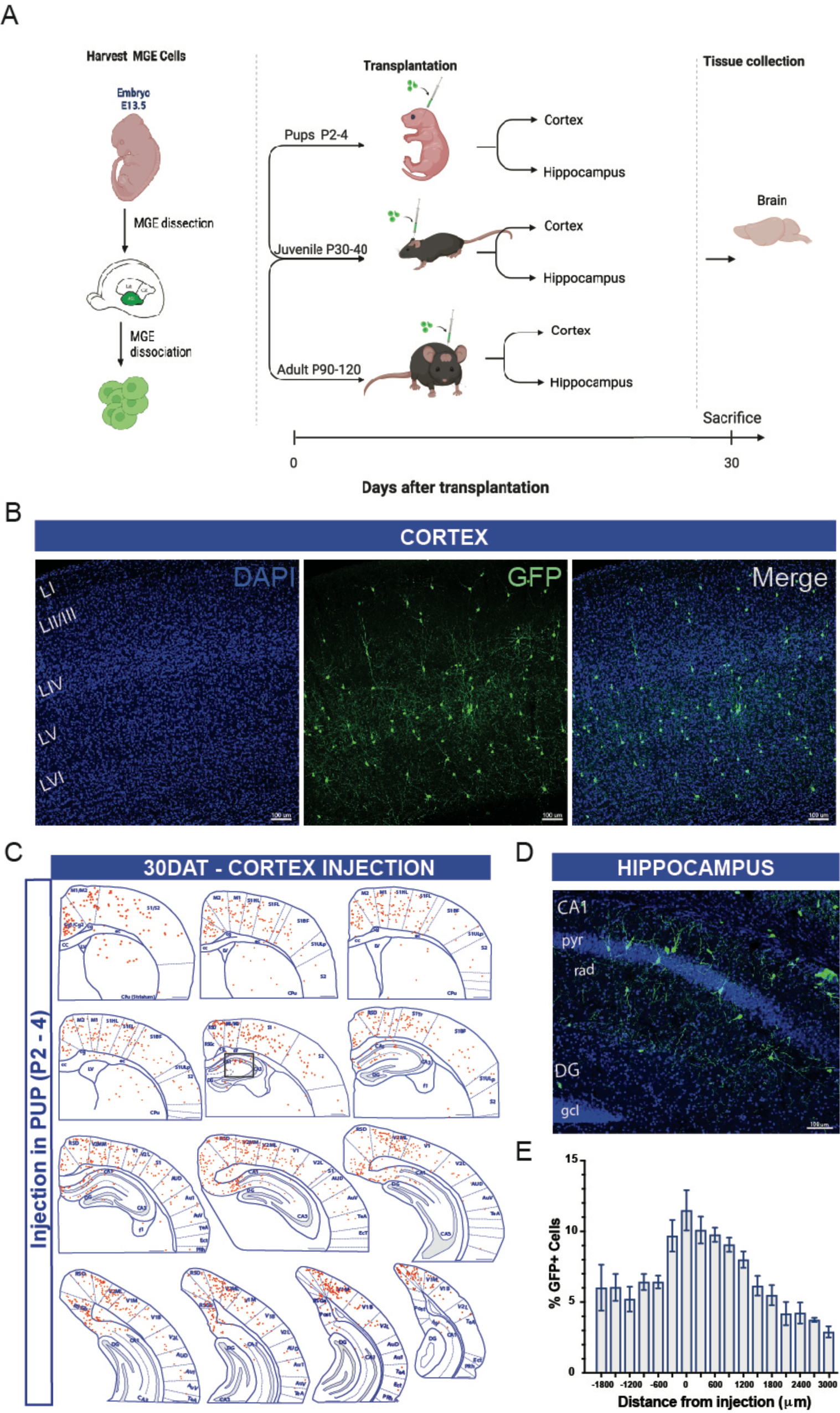
Timeline and distribution of transplanted MGE progenitors in pup cortex. (A) Schematic of the overall experimental design. (B) Coronal section of a pup recipient mouse (30 DAT) labeled for transplanted GFP+ neurons (green). Transplanted cells widely dispersed across cortex. (C) Schematic at 30 DAT showing distribution of GFP+ cells (red) across brain slices in the antero-posterior axis (300 μm apart) after transplantation in pup cortex. Region divisions were adapted from the Paxinos atlas. (D) Higher magnification of the hippocampal region outlined in B. (E) Plot showing the distribution of GFP+ cells in the antero-posterior axis 30 DAT cortex injection in pup (P2-4) mice; n = 8 brains. DAT, days after transplantation. Data represented as mean ± SEM.

### MGE progenitors transplanted into cortex

To investigate the influence of host brain age on survival, migration and maturation, we harvested MGE progenitor cells from E13.5 GFP-labeled donor embryos and transplanted into neocortex of neonatal (P2-P4), juvenile (P30-40) and adult (P60-150) mice. At 30 days post-transplantation (DAT), we noted a significant difference in cell survival across the three age groups with higher GFP+ cell survival rate following transplantation into neonatal host brain compared to juvenile or adult mice (P2-4, 17.2 ± 2.5%, n = 8 mice; P30-40, 0.8 ± 0.1%, n = 6 mice; P60-150, 0.7 ± 0.1%, n = 5 mice; p <0.001, one-way ANOVA). We also found, a significant difference in migration capacity away from the injection site. At 30 days after a single cortical injection of MGE-GFP progenitors in neonates (n = 8), GFP+ cells showed extensive distribution along the antero-posterior, medial-lateral and dorsal-ventral axis (Fig. 1B-E). In the anterior-posterior direction, cells migrated up to 3000 µm from the injection site in both directions covering the entire antero-posterior cortical structures (up to anterior cingulate and visual cortex in anterior and posterior axis, respectively). Along the dorsal-ventral axis, in addition to neocortex, MGE-GFP cells reached into striatum, dorsal subiculum and in some cases CA1 regions of hippocampus (Fig. 1D). Across the medial-lateral extent of neocortex, MGE-GFP cells reached lateral cortical areas such as auditory cortex. In juvenile and adult host brain, even though transplanted MGE-GFP progenitor cells migrated away from the injection site, the distance reached in antero-posterior axis was more limited (up to 600 µm) (Figs. 2A-B). Total AP cortical distances covered were 3825 ± 239 µm (neonate), 850 ± 92 µm (juvenile) and 900 ± 95 µm (adult), respectively (p <0.001, one-way ANOVA).

**Figure 2:**
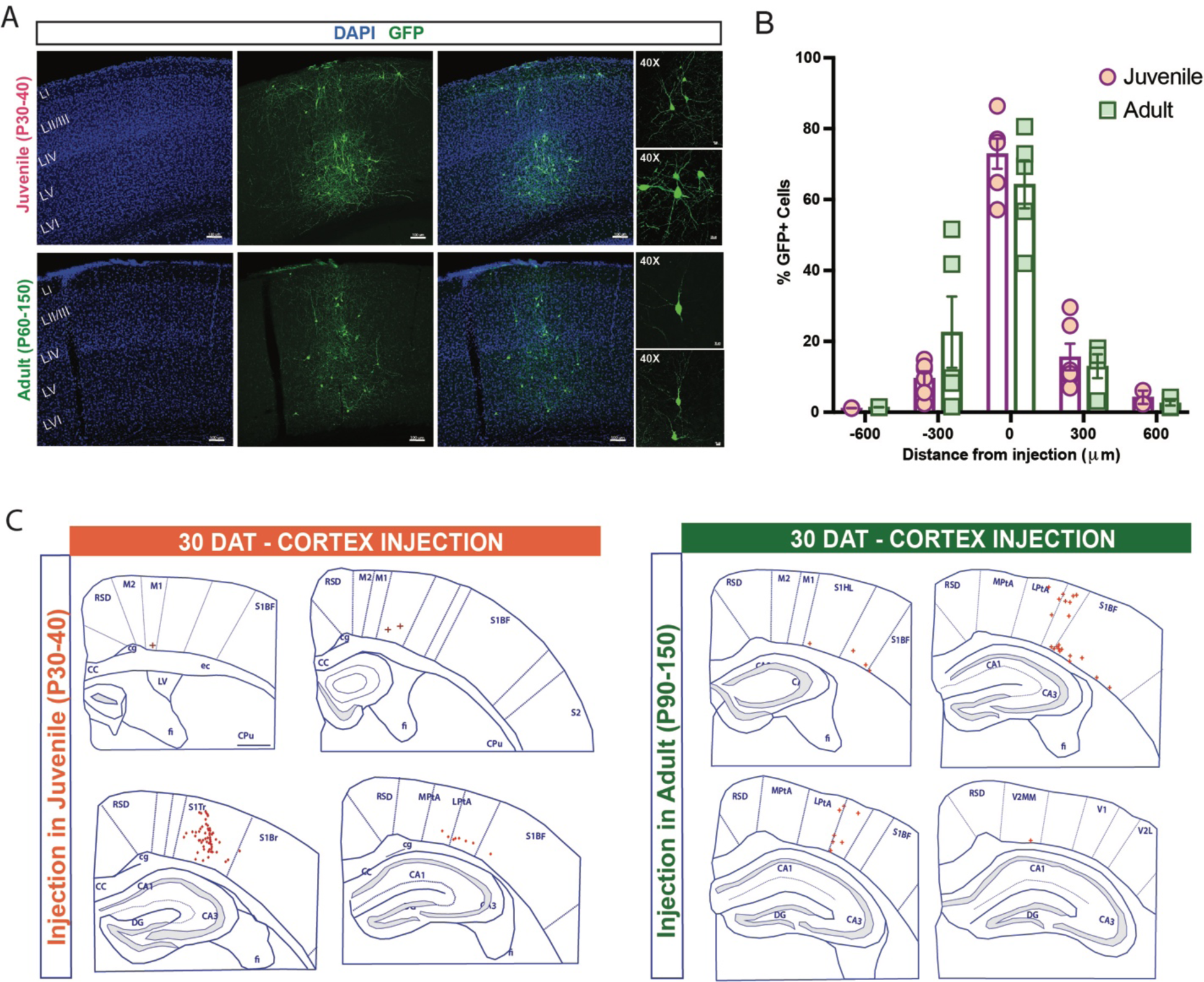
Transplanted MGE progenitors in juvenile and adult cortex. (A) Representative images from 30 DAT brains receiving MGE progenitor cells in the cortical area at P30 (juvenile, top) and P90 (adult, bottom). (B) Plot showing the distribution of GFP+ cells in the antero-posterior axis from the injection site in juvenile (P30-40) mice; n = 6 brains and adult (P60-150) mice, n = 5 brains. (C) Schematic at 30 DAT showing distribution of GFP+ cells (red) across brain slices in the antero-posterior axis (300 μm apart) after transplantation in juvenile (left panel) and adult (right panel) cortex. Region divisions were adapted from the Paxinos atlas. Data represented as mean ± SEM.

Next, to assess laminar distribution of MGE-derived GFP+ cells within specific cortical sub-layers of host brain, we divided the radial axis of cortex into ten equal bins and assigned each GFP+ cell to a single bin (see Methods). In the neonatal (P2-P4) transplantation group, GFP+ cells were distributed across the entire neocortex from Layer II to VI (Fig. 3) and no cells were identified in Layer I (which corresponds to bin 1), matching endogenous MGE-derived interneurons cortical distribution ^30–32^. In juvenile (P30-40) and adult (P60-150) transplantation groups, we observed a similar distribution of GFP+ cells across layers. However, an increased number of transplant-derived cells were observed in deep cortical layers (bin 10) near injection sites in adult compared to pup and juvenile mice (Fig. 3A-B). Also, along medial-lateral axis, we found a significant increase in GFP+ cells that migrate laterally from the injection site after transplantation in neonatal host brain compared to juvenile and adult (P2-4, 5229 ± 80.1 mm, n = 8 mice; P30-40, 970 ± 66.6 mm, n = 6 mice; P60-150, 994 ± 62 mm, n = 5 mice; p <0.001, one-way ANOVA) (Fig. 3C). Interestingly, in juvenile and adult brains, transplanted-derived cells appeared to have columnar distribution while in pup brains, they were broadly distributed.

**Figure 3:**
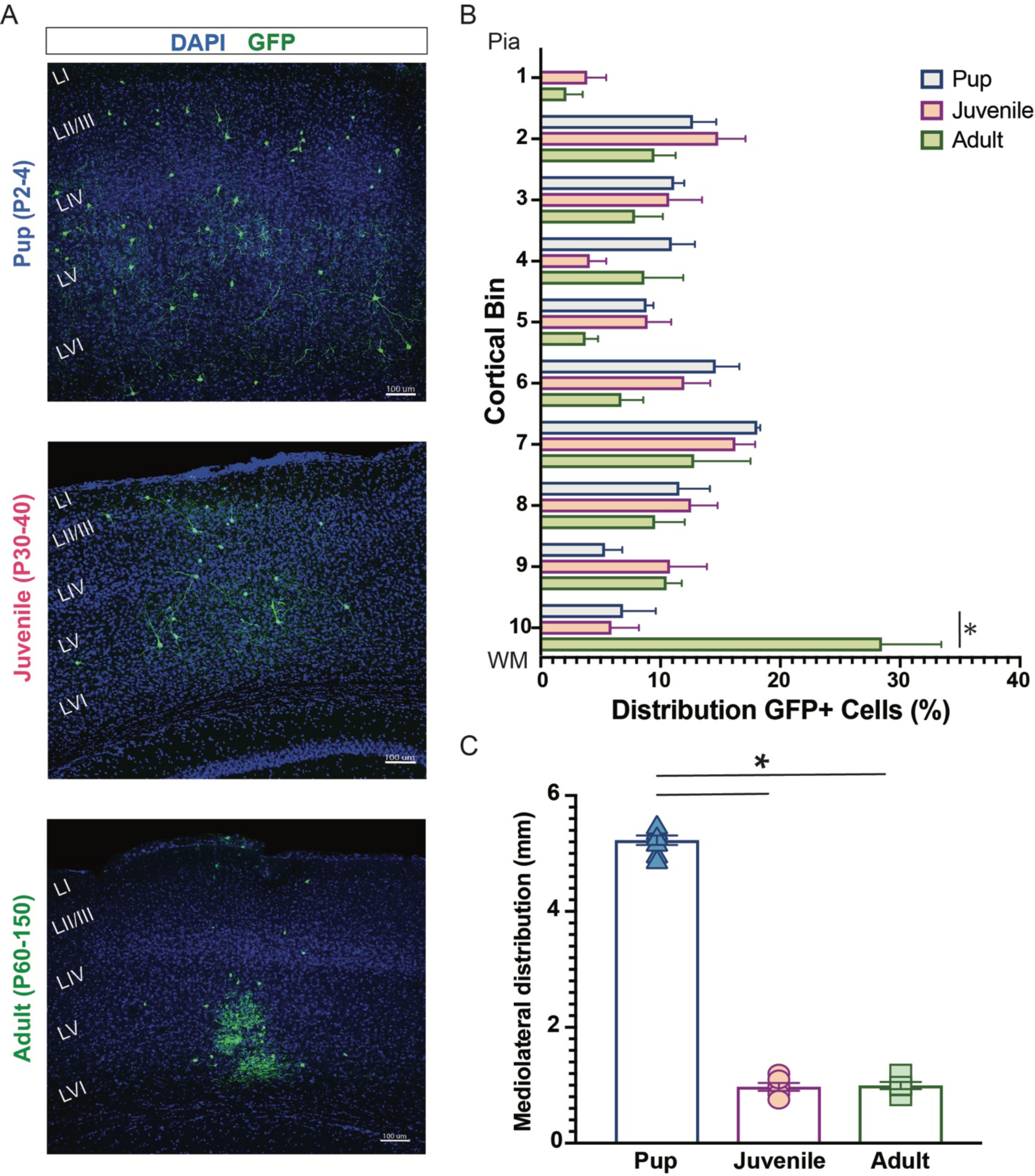
Distribution of MGE-derived cells following transplantation in cortex. (A) Representative coronal sections at 30 DAT after pup (top), juvenile (middle) and adult (bottom) transplantation. (B) Plot showing the distribution of MGE-derived cells along rostro-caudal axis divided in 10 bins from pia to white matter (WM). Note: comparable distribution across different age groups except for an increased number of GFP+ cells close to injection site in adult (bin 10). (C) Plot of the mediolateral distribution of cells across age groups. Note a highly mediolateral migration after pup transplantation. DAT, days after transplantation. Data represented as mean ± SEM; One-way ANOVA * p < 0.05.

To evaluate whether host brain environment influences maturation of MGE-derived GFP+ cells, we performed a series of immunohistochemical studies using antibodies recognizing neurons (NeuN), interneurons (PV and SST) and glia (GFAP, Olig2 and CC1). In the neonatal (P2-P4) transplantation group, GFP+ cells were primarily NeuN-positive and co-labeled with PV or SST, as expected for MGE-lineage interneurons ^15,38^. GFP+ cells did not co-label with glial markers (Figs. 4A-B). In the juvenile (P30-40) transplantation group, a similar distribution of NeuN+, PV+ and SST+ co-labeled MGE-GFP cells was observed with few to no GFP+ cells co-labeled with GFAP, Olig2 or CC1 excluding the injection site. In the adult (P60-150) transplantation group, we noted an overall decrease in NeuN+ neurons (Fig. 4B) and, specifically PV+ interneurons, compared to younger transplantation groups (P2-4, PV: 28 ± 2.3%; P30-40, PV: 16.6 ± 4%; P60-150, PV: 10.5 ± 2%, p <0.01, one-way ANOVA). However, GFP+ cells co-expressed a similar proportion of SST (P2-4, SST: 33 ± 1.8%; P30-40, SST: 34.7 ± 3.4%; P60-150, SST: 30.9 ± 4.2%, p=0.7, one-way ANOVA). Interestingly, in juvenile and adult groups, we observed a few GFP+ cells co-expressing GFAP concentrated at the injection sites surrounded by an increased number of endogenous GFAP+ cells surrounding the injection site compared to adjacent cortical area as shown in Figure 5A. At distances of 300 um away from the injection site, MGE-GFP cells differentiated to cells with a neuronal morphology expressing neuron- and interneuron-specific markers (Fig. 5B). Taken together, these findings suggest neonatal cortex is a more permissible environment for transplantation of MGE progenitor cells compared to later stages of development.

**Figure 4:**
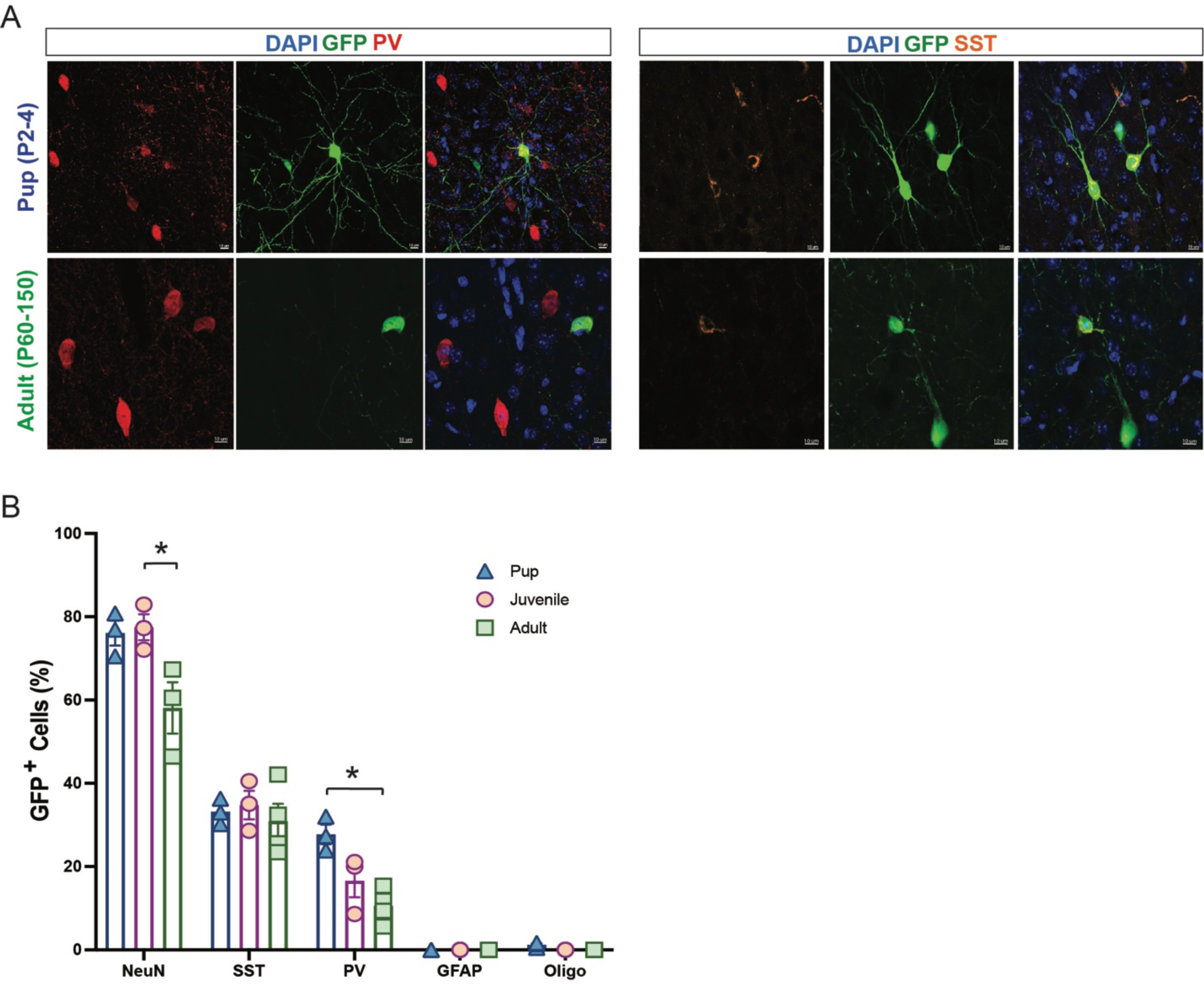
Maturation profile for MGE-derived interneurons transplanted in cortex. (A) Representative magnified cell images showing co-localization between GFP and PV (left side) and SST (right side) at 30 DAT after pup and adult transplantation. (B) Plot showing the quantification of marker expression in GFP-labeled cells (n = 3 mice per marker) across age groups. Note a decreased number of GFP co-expressing PV cells in juvenile and adult group. Data represented as mean ± SEM; One-way ANOVA * p < 0.05. DAT, days after transplantation. NeuN, neuronal nuclei; SST, somatostatin; PV, parvalbumin; GFAP, glial fibrillary acid protein; Oligo, oligodendrocyte.

**Figure 5:**
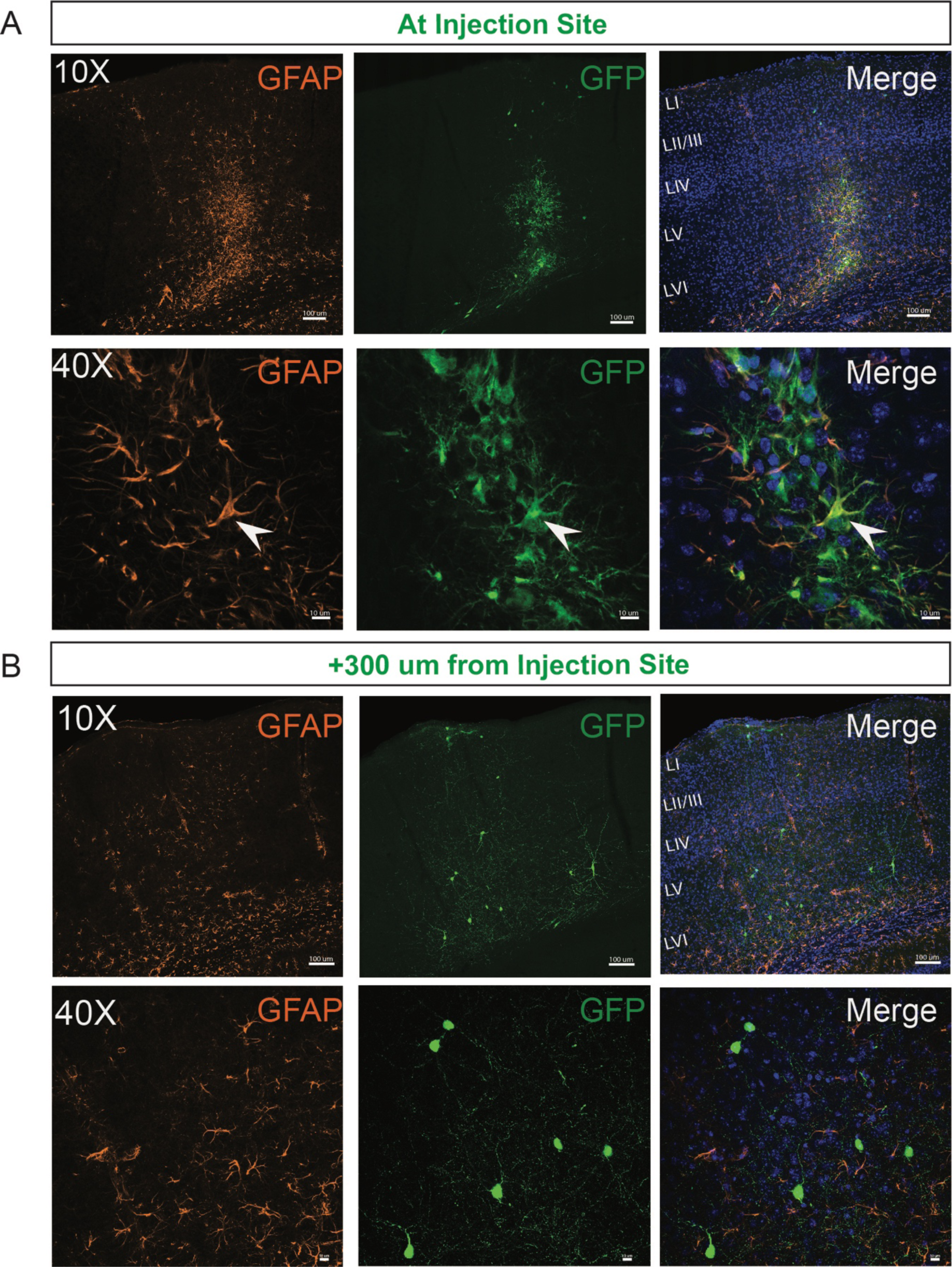
Endogenous astrocytes expression at the injection site. (A) Representative images collected at the injection site of an animal at 30 DAT. Note the presence of endogenous astrocytes (GFAP) around cortical injection site and a few GFP+ astrocytes (white arrow). (B) Representative images collected 300 μm away from the injection site from the same animal. GFP+ cells show mature neuronal morphology. DAT, days after transplantation. GFAP, glial fibrillary acid protein.

### MGE progenitors transplanted into hippocampus

To investigate if local host brain microcircuits influence survival, migration or maturation of transplanted MGE progenitor cells, we also targeted hippocampus. This complex brain region has a unique anatomical structure and is widely implicated in pathophysiology of neurological diseases^39^. MGE progenitors from E13.5 GFP-labeled donor embryos were transplanted into hippocampus of neonatal (P2-P4), juvenile (P30-40) and adult (P60-150) mice. At 30 DAT, we noted a significant difference in cell survival across the three age groups. A higher hippocampal GFP+ cell survival rate was noted in pup compared to juvenile and adult mice (P2-4, 8.7 ± 0.6%, n = 6 mice; P30-40, 0.98 ± 0.2%, n = 6 mice; P60-150, 0.7 ± 0.08%, n = 7 mice; p <0.001, one-way ANOVA).

Migration away from the injection site was also different across the three age groups. In particular, after a single injection of MGE progenitor cells in pup hippocampus, transplanted cells distributed widely across the entire antero-posterior axis of the hippocampus (up to 2400 μm from the injection site) with GFP+ cells spread into dorsal and ventral hippocampus as shown in Figure 6A-C. In some cases (4 out of 6 mice), a few GFP+ transplanted cells migrated and spread into cortical areas and transition subiculum (Fig. 6B); note the average GFP+ survival rate outside hippocampus was 5.9 ± 1.5% (n = 6 mice). Migration in juvenile and adult mice (Fig. 7A) was more limited reaching up to 900 µm from the injection site (Fig. 7B).

**Figure 6:**
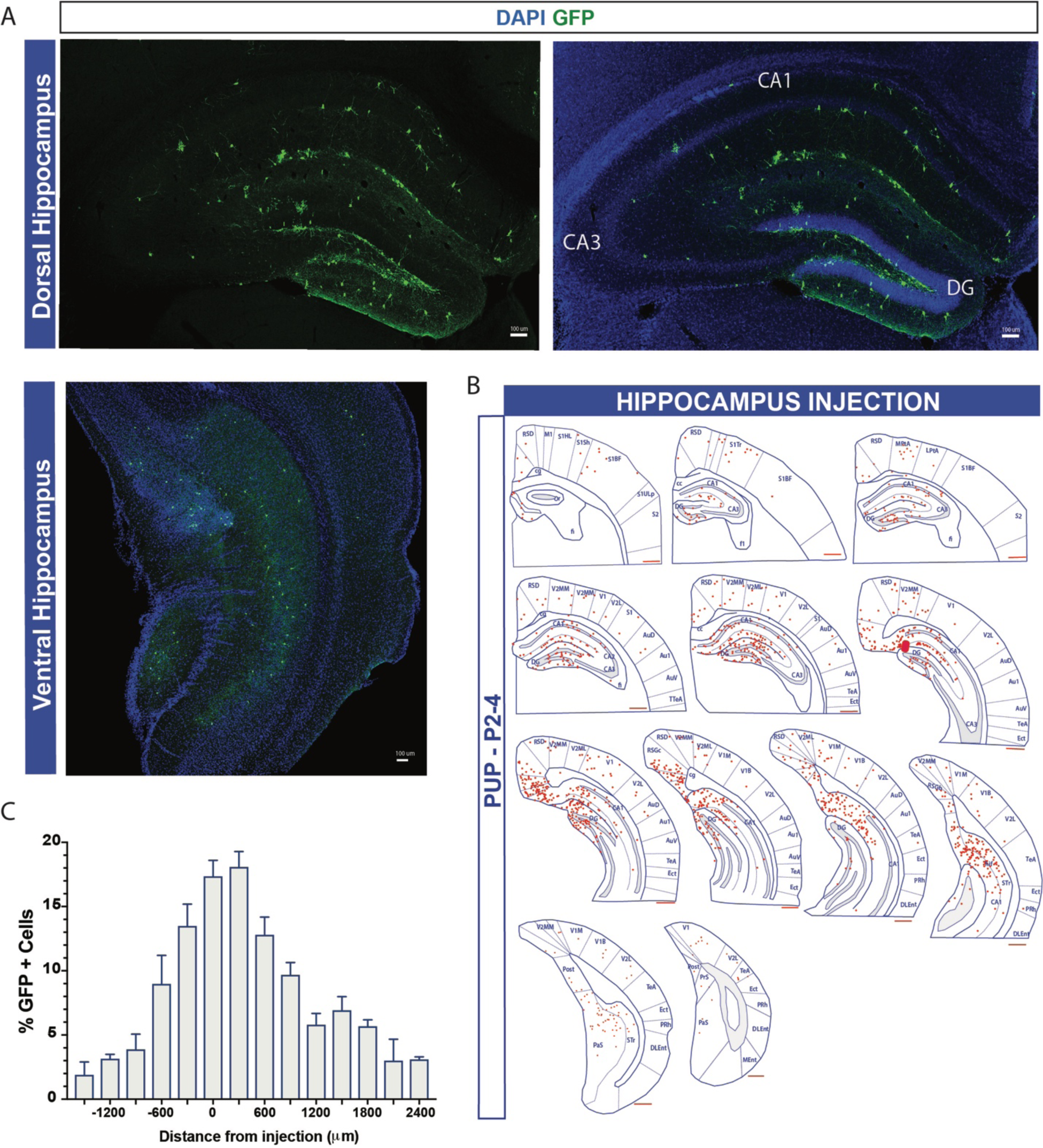
Distribution of MGE-derived cells following transplantation in hippocampus. (A) Representative images of dorsal and ventral hippocampus sections with GFP+ cells. (B) Schematic of the distribution of GFP+ cells (marked in red) section 300 μm apart. Region divisions were adapted from the Paxinos atlas. (B) Percentage of GFP+ cells across the antero-posterior axis (6 mice, from 3 or more experiments). (D) Plot showing the quantification of the total number of GFP+ in hippocampus at 30 DAT of a single injection. DAT, days after transplantation. Data represented as mean ± SEM.

**Figure 7:**
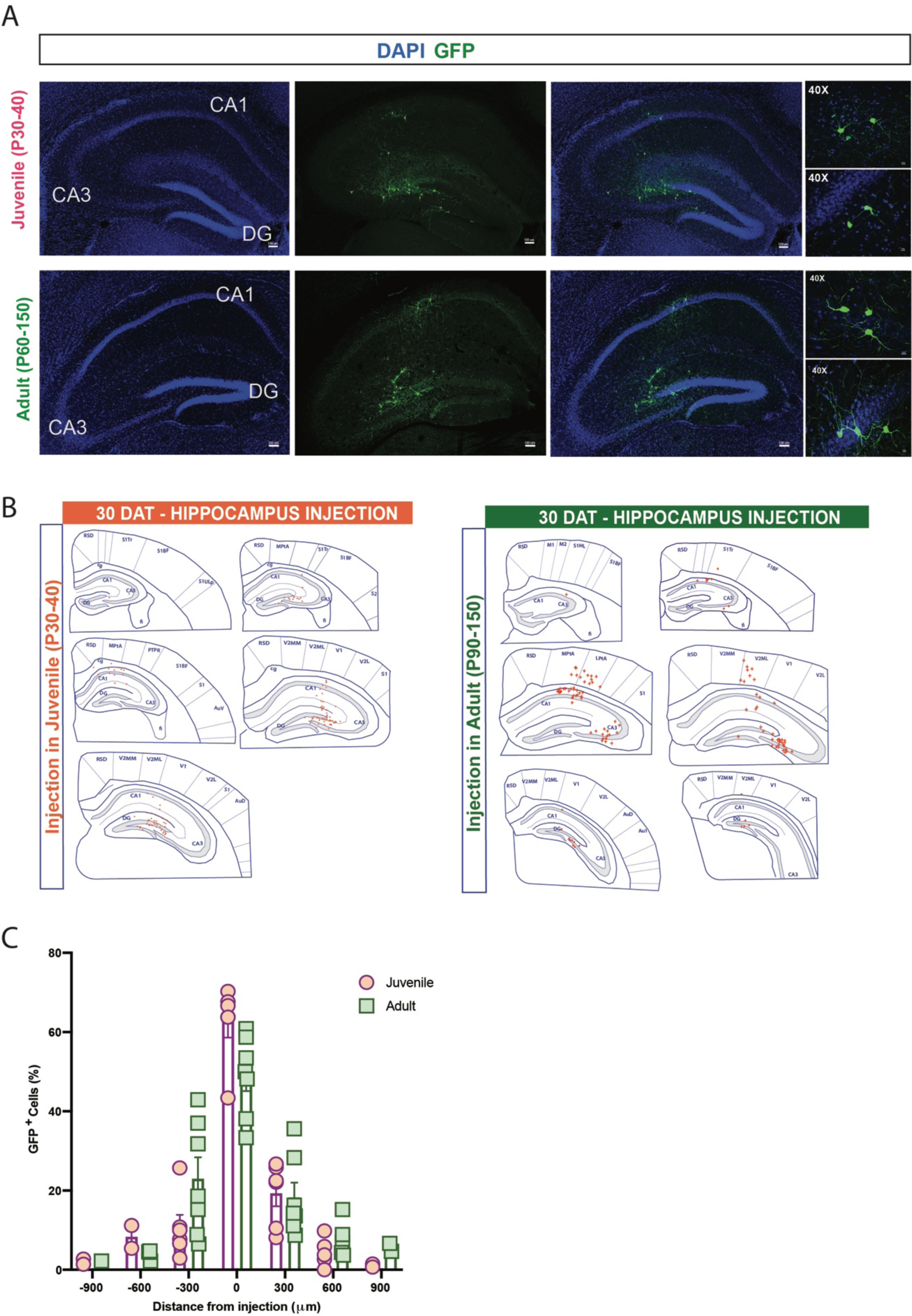
Transplanted MGE progenitors in juvenile and adult hippocampus. (A) Representative coronal section of hippocampus from juvenile (top) and adult (bottom) host brain at 30 DAT showing surviving GFP+ cells (green). Right panels show higher magnification of transplanted cells with interneuronal-like morphology. (B) Schematic at 30 DAT showing distribution of GFP+ cells (red) across brain slices in the antero-posterior axis (300 μm apart) after transplantation in juvenile (left panel) and adult (right panel) hippocampus. Region divisions were adapted from the Paxinos atlas. (C) Plot of the percentage of GFP+ cells across the antero-posterior axis (6 mice, from 3 or more experiments). DAT, days after transplantation. Data represented as mean ± SEM.

In all age groups (pup, juvenile and adult), transplant-derived cell distribution within hippocampus was not uniform and an increased number of GFP+ cells were observed in specific subregions. For example, at 30 DAT in hippocampal DG of pups, GFP+ cells were primarily found in DG and only few cells in CA1 and CA3 subregions (74, 21.4 and 4.5%, respectively; p <0.001, one-way ANOVA) (Fig. 8A). In adult mice, cells transplanted in area CA3 migrated mostly within CA3 subregions (70%) and only few cells reached CA1 or DG (Fig. 8B). To note, in one case we observed an extensive migration of GFP+ cells in DG after injection in CA3 stratum radiatum. Those results highlight that in addition to a host brain age effect, within hippocampus, targeted subregions also influence migration of transplanted MGE progenitor cells.

**Figure 8:**
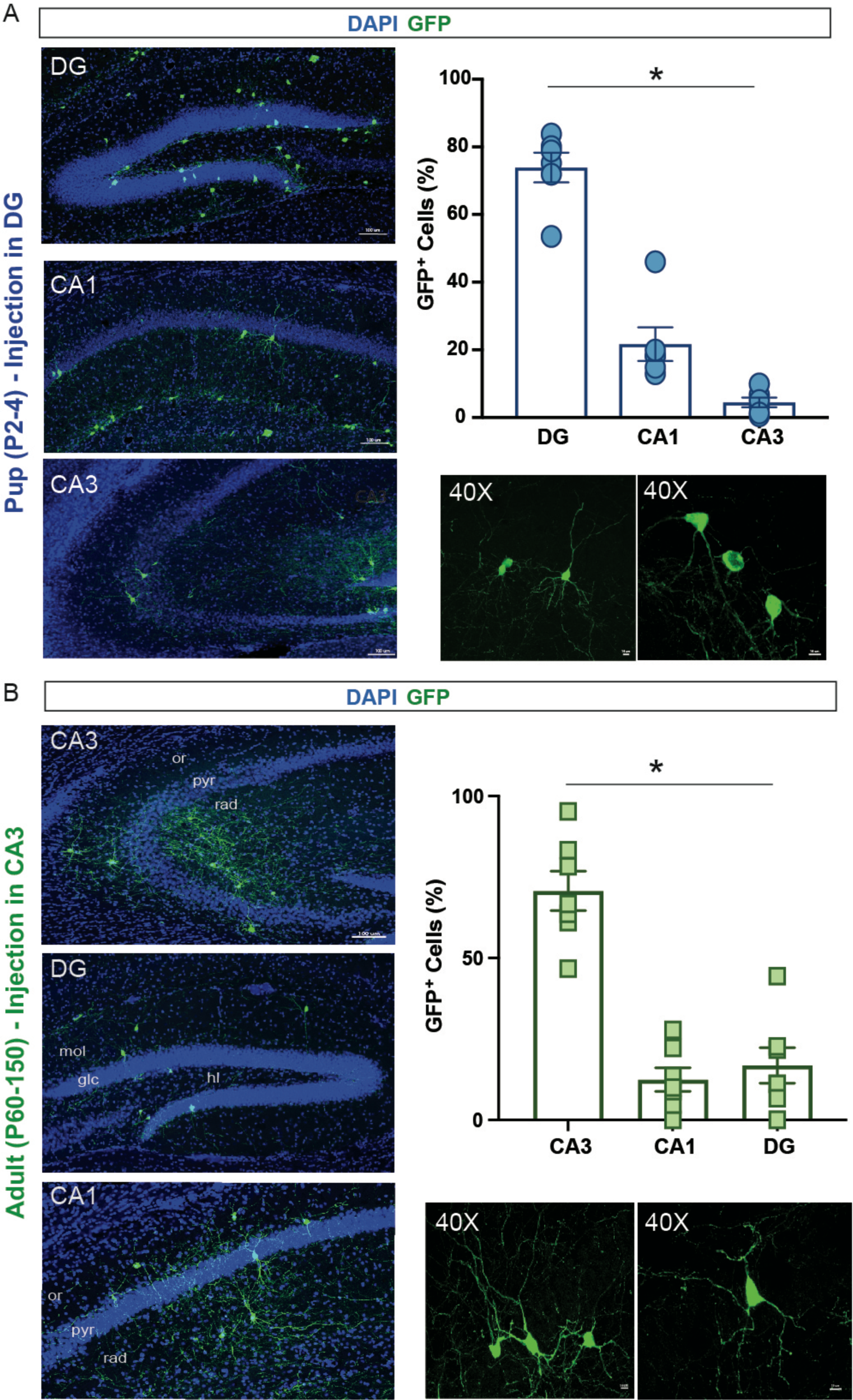
Distribution of transplant-derived MGE-GFP cells within hippocampal subfields. (A) Representative hippocampal sections of the subfield CA1, CA3 and DG at 30 DAT after pup injections. Plot showing the quantification of GFP+ cells across the subfields of hippocampus (6 mice from 3 or more experiments). Note increased number of cells in dentate gyrus compare to CA1 and CA3. (B) Representative hippocampal sections of the subfield CA1, CA3 and DG at 30 DAT showing the distribution of GFP+ cells after adult injections. Quantification of GFP+ cells across the subfields of the hippocampus (6 mice from 3 or more experiments). Note the increased number of cells in CA3 compared to CA1 and DG. Higher resolution examples of GFP+ cells imaged under 40X objective showing a mature neuronal morphology are shown at bottom. DAT, days after transplantation; GFAP, glial fibrillary acid protein. Data represented as mean ± SEM; one-way ANOVA * p < 0.05.

Next, we evaluated whether hippocampal host brain environment influences differentiation of MGE-derived cells. No differences were observed in GFP+ cells co-expressing NeuN (P2-4, NeuN: 49.5 ± 1.6%; P30-40, NeuN: 69.1 ± 1.9%; P60-150, NeuN: 70.3 ± 9.1%, p>0.05, one-way ANOVA), PV (P2-4, PV: 9.2 ± 3.1%; P30-40, PV: 6.1 ± 2.4%; P60-150, PV: 3.4 ± 1.8%, p>0.05, one-way ANOVA), Oligo (P2-4, Oligo: 1.6 ± 0.5%; P30-40, Oligo: 6.4 ± 4%; P60-150, Oligo: 0%, p>0.05, one-way ANOVA) and CC1 (P2-4, CC1: 0.9 ± 0.7%; P30-40, CC1: 0.9 ± 0.9%; P60-150, CC1: 0%, p>0.05, one-way ANOVA) across the three age groups (Fig. 9). However, we observed reduced GFP+ cells co-expressing SST in pup compared to juvenile and adult (P2-4, SST: 10.7 ± 1.7%; P30-40, SST: 38 ± 2.7%; P60-150, SST: 36.2 ± 5.4%, p<0.01, one-way ANOVA, Fig 9B). Surprisingly, around 13% of transplanted MGE progenitor cells in pup hippocampus (DG area), co-expressed GFAP at 30 DAT (Fig. 10A) while no GFAP+ cells were observed in juvenile or adult mice. GFP+ co-expressing GFAP cells migrated along the antero-posterior axis up to 1500 μm from the injection site (Fig. 10B) and were only located within DG. These cells exhibit morphology, size and branch orientation similar to molecular astrocytes recently described by Karp et al.^40^ and distributed across dentate gyrus layers (ML, GZ, SGZ, hilus) similar to endogenous astrocytes (Fig 10C).

**Figure 9:**
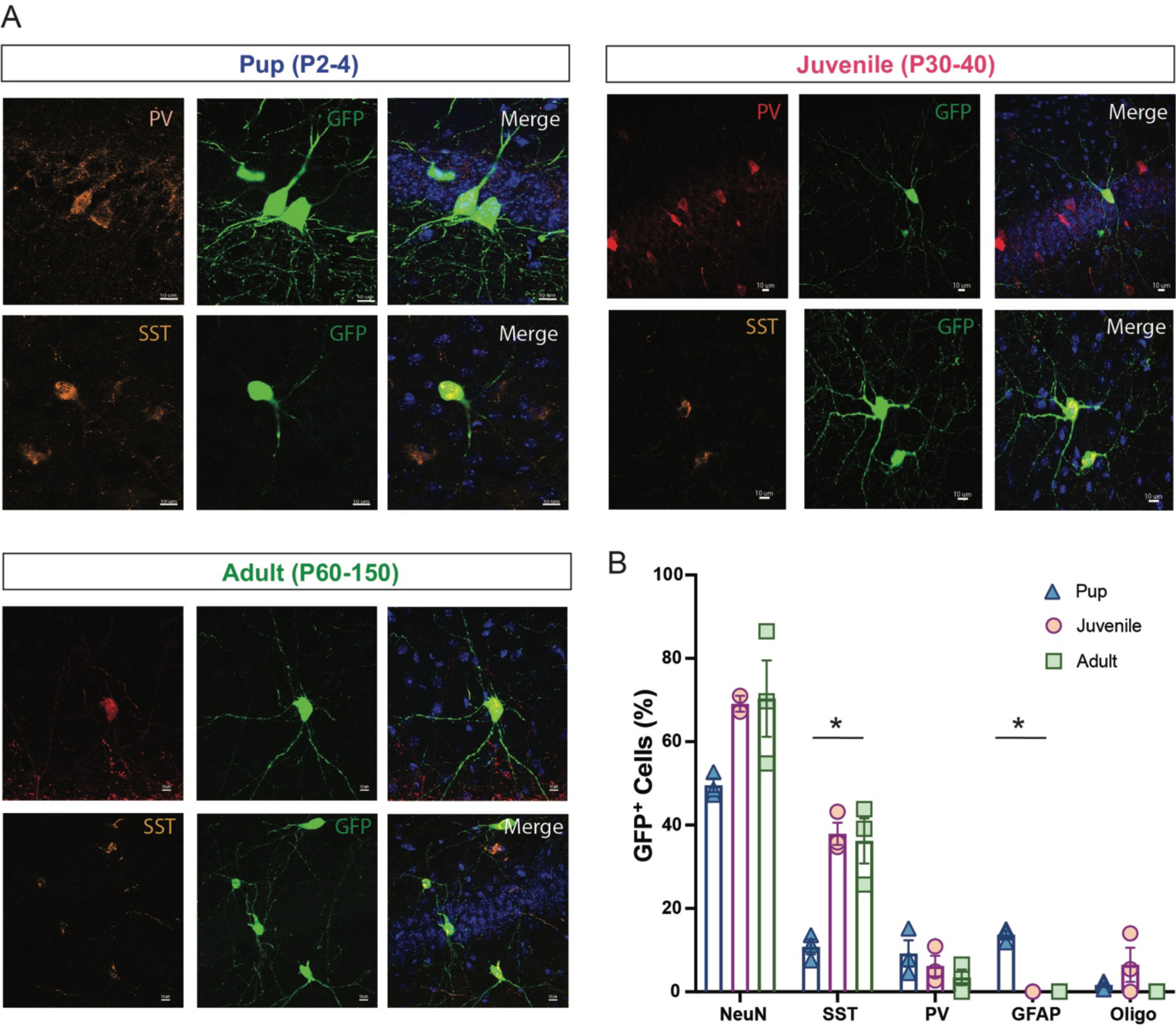
Maturation profile for MGE-derived interneurons transplanted in hippocampus. (A) Representative magnified cell images showing GFP+ cells co-expressing PV and SST at 30 DAT in pup, juvenile and adult hippocampus. (B) Plot showing the quantification of cells co-expressing GFP and NeuN, SST, PV, GFAP, Oligo in hippocampus at 30 DAT in for pup, juvenile and adult groups (n = 3-4 mice per marker). Note reduced GFP+ cells co-expressing SST and increased GFP+ co-expressing GFAP cells in pup group. DAT, days after transplantation. Data represented as mean ± SEM; one-way ANOVA * p < 0.05. Abbreviations as in Figure 4.

**Figure 10:**
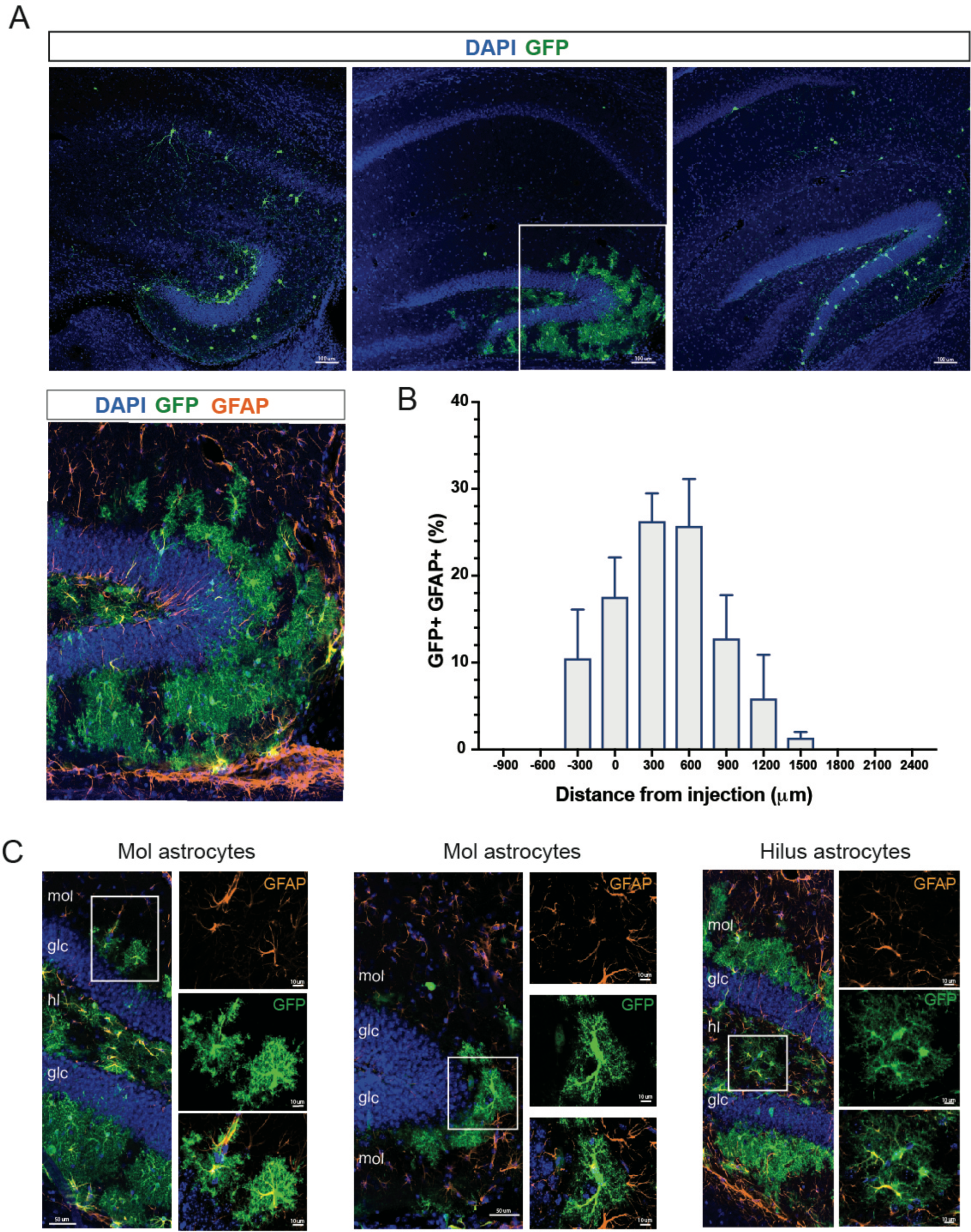
Morphologically distinct astrocytes across DG layers. (A) Representative coronal images of GFP+ cells in hippocampus at different anteroposterior levels following transplantation at P2-4. Higher resolution image of the dentate gyrus showing distribution of GFP+ co-expressing GFAP across dentate gyrus layers (bottom, left). (B) Plot showing the distribution of GFP+ co-expressing GFAP across the antero-posterior axis. (C) Representative images of GFP+ co-expressing GFAP cell morphologies at the molecular layer (left, middle) and hilus (right). Higher magnification (40x objective; white box) images cells are shown at right. DAT, days after transplantation. Data represented as mean ± SEM.

## Discussion

Our results suggest that host brain environment is responsible for regulating survival, migration and maturation of cells derived from transplantation of embryonic MGE progenitors. Although previous work suggested that interneuron survival following transplantation is guided by expression of an intrinsic maturational program, rather than developmental state of the host brain itself ^41^, our data effectively demonstrates robust survival and extensive migration when MGE progenitors are transplanted early (but not later) in development or into cortex (compared to hippocampus). Maturation profile of MGE-derived interneurons is also regulated by developmental state and brain region as we observed a two-fold greater percentage of SST-positive interneurons following hippocampal transplantation into juvenile or adult brain (compared to neonates); whereas the percentage of PV-positive interneurons decreased nearly two-fold with age following cortical transplantation into adult brain (compared to neonates). Unexpectedly, we also found sub-regions of pup hippocampus – namely the dentate gyrus – where transplant-derived MGE progenitors locally differentiate into astrocytes. Considering the emerging therapeutic potential for transplanted embryonic MGE progenitors in a variety of neurological disorders ^10,11,13,21,25,42^, these results have important clinical implications.

There are several mechanisms that could regulate survival, spatial distribution and maturation of MGE-derived interneuron subtypes following transplantation. First, activity-dependent mechanisms - such as excitatory pyramidal cell activity - may serve as a critical regulator of interneuron development establishing age- and region-specific permissive corridors for MGE progenitors ^43–45^. In murine neocortex, for example, spontaneous patterns of activity present at birth ^46–48^ could facilitate the widespread anterior-to-posterior migration of transplanted MGE progenitors seen here (Fig. 1). Second, precursors for MGE-derived interneuron subtypes could be subject to programmed cell death during neurodevelopment. This process is regulated by region-specific expression of environmental factors like *pcdh* ^34,35,49^, *c-Maf* ^50^ or *PTEN* ^44^ which facilitate postnatal survival and/or maturation of PV+ and SST+ interneurons. Given that these are developmentally regulated expression patterns it is not surprising that juvenile and adult brains were less receptive to integration of transplanted MGE progenitors than mouse pups or the differences in PV/SST ratios seen here (Fig 4).

In our study, few GFP+ cells co-expressed classical oligodendrocyte ^51–53^ or astrocyte ^54^ markers around the injection site following MGE transplantation in juvenile/adults mice, as previously reported ^10,18,33,55^. Surprisingly, following MGE transplantation into pups’ hippocampus, a significant population of GFP+ cells co-expressing GFAP was distributed around the DG layers showing an endogenous astrocyte-like morphology. Numerous studies have shown that glia cells can also originate from Nkx2.1-expressing precursor of embryonic MGE ^56^ and after migrating in the cortex, they transiently participate in cortical neurodevelopment to be later lost and replaced by other glia cell populations^57^. Furthermore, earlier studies showed the absence of proliferative markers following transplantation of embryonic MGE progenitors ^22,58^ and while unlikely, the presence of proliferating neural cells cannot be fully excluded. Future studies focused on additional markers and lineage analysis will be necessary to identify factor(s) driving astrocyte survival following transplantation into the DG microenvironment.

In our hands, MGE transplantation at postnatal day 2 has consistently been shown to result in widespread migration, in all directions, within host brain parenchyma ^18,30,26^ transplanted MGE progenitor cells also show clear morphologies consistent with migrating cells ^18^. Our results confirm and extend these reports. While the possibility that some MGE-GFP cells distributed to distant sites via the meninges or leakage into the cerebral spinal fluid cannot be entirely excluded given the small size of the mouse brain ^59^, the migratory profiles and properties of MGE-derived cells both *in vivo* and *in vitro* do not support this conclusion. However, it is interesting that in wild-type juvenile or adult mice the migratory ability was more limited. The different molecular, cellular, extracellular profiles in the adult versus pup environment ^60,61^ may explain this reduced migration in mature brain compared to early developmental stages. Such difference, while it might seem a limitation of the technique, highlights instead the translational applicability of this approach given the ability to modify host circuits with regional and neuronal specificity in mature brain compared to the broad, non-specific circuit changes after pups’ transplantation. Indeed, adult MGE transplantation has already proven an effective disease-modifying therapy in adult mouse models representing acquired epilepsy or traumatic brain injury ^10,23,12^.

It is worth noting that a significant proportion of individuals with epilepsy, autism spectrum disorders or schizophrenia are considered “inter-neuropathies”, define by frank reductions or dysfunction in PV+ and/or SST+ interneurons ^3,62–64^. As these conditions present at different stages of development, and as interneuron deficits can be localized to cortical or hippocampal brain regions, a better understanding of the influence of local environment on transplanted MGE progenitors is necessary. Indeed, several studies using embryonic MGE progenitors ^65^ or other embryonic cell sources ^66,67^ for transplantation also support our findings that local environmental factors influence the integration of these cells in the host brain. Understanding advantages and limitations of where (and when) MGE progenitors integrate following transplantation is critical to establishing effective targeted cell therapies for patients suffering from a variety of neurological disorders.

## Author contributions

R.P and S.C.B. designed research; R.P, T.V. C.H. conducted the research R.P and T.V. analyzed data; R.P and S.C.B. wrote the paper.

